# Response to photic stimulation as a measure of cortical excitability in epilepsy patients

**DOI:** 10.1101/2023.04.23.537976

**Authors:** Michaela J. Vranic-Peters, Patrick O’brien, Udaya Seneviratne, Ashley Reynolds, Alan Lai, David Grayden, Mark Cook, Andre Peterson

## Abstract

Studying states and state transitions in the brain is challenging due to nonlinear, complex dynamics. In this research, we analyse the brain’s response to non-invasive perturbations. Perturbation techniques offer a powerful method for studying complex dynamics, though their translation to human brain data is under-explored. This method involves applying small inputs, in this case via photic stimulation, to a system and measuring its response. Sensitivity to perturbations can forewarn a state transition. Therefore, biomarkers of the brain’s perturbation response or ‘cortical excitability’ could be used to indicate seizure transitions. However, perturbing the brain often involves invasive intracranial surgeries or expensive equipment such as transcranial magnetic stimulation (TMS) which is only accessible to a minority of patient groups, or animal model studies. Photic stimulation is a widely used diagnostic technique in epilepsy that can be used as a non-invasive perturbation paradigm to probe brain dynamics during routine electroencephalography (EEG) studies in humans. This involves changing the frequency of strobing light, sometimes triggering a photo-paroxysmal response (PPR), which is an electrographic event that can be studied as a state transition to a seizure state. We investigate alterations in the response to these perturbations in patients with genetic generalised epilepsy (GGE), with (*n* = 10) and without (*n* = 10) PPR, and patients with psychogenic non-epileptic seizures (PNES; *n* = 10), compared to resting controls (*n* = 10). Metrics of EEG time-series data were evaluated as biomarkers of the perturbation response including variance, autocorrelation, and phase-based synchrony measures. We observed considerable differences in all group biomarker distributions during stimulation compared to controls. In particular, variance and autocorrelation demonstrated greater changes in epochs close to PPR transitions compared to earlier stimulation epochs. Comparison of PPR and spontaneous seizure morphology found them indistinguishable, suggesting PPR is a valid proxy for seizure dynamics. Also, as expected, posterior channels demonstrated the greatest change in synchrony measures, possibly reflecting underlying PPR pathophysiologic mechanisms. We clearly demonstrate observable changes at a group level in cortical excitability in epilepsy patients as a response to perturbation in EEG data. Our work re-frames photic stimulation as a non-invasive perturbation paradigm capable of inducing measurable changes to brain dynamics.

## 1 Introduction

Understanding brain function is challenging due to its’ nonlinear, complex dynamics. Methods from statistical physics and dynamical systems theory have been successful in characterising other complex, nonlinear systems, including ecological population changes [1], weather [2], and economics [3]. These theories have also been successfully applied to the study of the brain, usually in animal models or using invasive procedures, including the study of brain state transitions in epilepsy patients [4; 5; 6; 7].

Epilepsy is a brain disease characterized by abnormal electrical brain activity and unprovoked recurrent seizures, that occurs in approximately 1% of the population [8]. These can be detected in electroencephalography (EEG), and the transition to seizure can be conceptualised as a critical transition [4; 9; 10]. Metrics that identify changes in system dynamics before these transitions, called ‘early warning signs’, have been observed in the EEG of epilepsy patients before seizures [11; 7]. These include increased variance, autocorrelation, and sensitivity to perturbation (a small external input) [12].

As a system approaches a state transition, it becomes more sensitive to perturbations, such that it is slower to return to equilibrium [12]. Since increased sensitivity to perturbation can forewarn a state transition, biomarkers of the brain’s perturbation response or ‘cortical excitability’ [13] could be used to track brain dynamics towards transitions to different states, including seizure states or other altered states of consciousness. Studying epilepsy in humans often involves trying to capture seizures during brain recordings, which can never be guaranteed and often requires long-term recordings. Perturbation methods have the benefit of reliably inducing responses of interest in real time. However, delivering perturbations to the brain is generally invasive in practice, requiring implantation of intracranial electrodes or other invasive devices. Animal studies in mice have successfully used electrical stimulation as a perturbation to track cortical excitability [14]. In humans, perturbation studies have used a subset of patients with focal epilepsy eligible for resection surgeries who are undergoing monitoring to confirm the seizure onset zone, where electrodes are placed strategically for that patient’s unique seizure onset zone [15; 16; 17; 18; 19]. Non-invasive perturbation is also possible through technologies such as transcranial magnetic stimulation (TMS) [20; 21; 22], although this technology is relatively expensive and the perturbation itself is arguably diffuse and spatially non-specific.

Photic stimulation is a widely used diagnostic technique [23] that could be used as a non-invasive perturbation paradigm to probe brain dynamics during routine EEG studies in humans. It involves changing the frequency of strobing light, sometimes triggering a photo-paroxysmal response (PPR), which is an electroencephalographic event visible on EEG. Generally, its primary clinical relevance is to diagnose photosensitive epilepsy, although we argue that it offers a safe, inexpensive research method for brain perturbation studies that is accessible to a much broader range of people. Furthermore, unlike stimulating the brain with electrodes or with TMS, the stimulation itself is input via the visual system of the brain, engaging with the brain’s functionality in a more natural endogenous way. In previous studies, markers of critical transitions have previously only been observed in patients with focal epilepsy; we aim to observe them during the generalised PPR response. Also, unlike previous studies, we aim to compare different epileptic populations in order to compare how photic stimulation effects these different groups, including patients with genetic generalised epilepsy (GGE) who are photosensitive and GGE patients who are not photosensitive, as well as patients with Psychogenic Non-Epileptic Seizures (PNES) and healthy controls.

Kalitzin [6] and Parra [24] found increased excitability in photosensitive individuals during stimulation using a measure of phase clustering called relative phase clustering index (rPCI). An extended study used rPCI to indicate greater probability of epileptic seizures [15]. In another study, higher visually evoked potential (VEP) amplitudes indicated greater responses to photic stimulation in individuals with photosensitive epilepsy compared to controls, especially in occipital areas [25]. In fact, individuals with photosensitive epilepsy exhibited stronger suppression of visual perception after TMS over the visual cortex, and had lower phosphene thresholds compared to healthy controls, indicative of increased excitability of the visual cortex.

Studies have used power spectrum measures to indicate changes in excitability. For instance, reduced inhibition of alpha rhythm generating networks was associated with photosensitive epilepsy [26], and stimulation via flashed chromatic gratings induced alpha desynchronisation in posterior electrodes [27]. Changes in power spectral density (PSD) in lower frequencies have been demonstrated to predict seizures from EEG recordings [28], which is consistent with expectations of increased autocorrelation and has a theoretical relationship with the power spectrum [29], such that we may expect increased low frequency power and long-range temporal associations. Therefore, increased spatial similarity could also be a candidate EWS, that could be measurable via synchrony measures like phase clustering and phase synchrony. This is supported by findings of increased phase synchronisation in the gamma frequency range in EEG, being associated with increased neuronal excitability in epilepsy patients [11].

We extend these analyses by using the PPR and photic stimulation from the perspective of dynamical systems theory. We study the transition to PPR as a critical transition, and expect to observe early warning signs prior to that transition.

As mentioned, we expect increased response to perturbation, variance, and autocorrelation in the brain. The goal of this research is to measure cortical excitability in epilepsy patients via a perturbation response. In this case, the perturbation response is non-invasive and in the form of photic stimulation, which is a commonly used investigation in routine EEG studies. We hypothesise that we can use photic stimulation as a perturbation method for observing increased cortical excitability before PPR. We expect to observe early warning signs of a PPR transition that are identifiable via changes to a particular metric’s distribution over time. We analyse distributions of statistical moments of the EEG timeseries as a data-driven method for inferring changes to system dynamics, because the underlying dynamics of brain activity are not observable. These include changes in variance and autocorrelation function widths of brain activity, as well as in phase-based synchrony measures. In particular, we expect the greatest increases to be demonstrated in the photosensitive GGE (PPR) group, just prior to a PPR, where they likely experience heightened cortical excitability.

## 2 Methods

### 2.1 Participants

The study was conducted using data obtained retrospectively from databases of the Department of Neurosciences, St. Vincent’s Hospital, Melbourne, over the period May 2012 - August 2022. This included selected EEG recordings with photic stimulation from 10 patients with genetic generalised epilepsy (GGE) and photoparoxysmal response (PPR) used in a previous study [30], 10 GGE patients without PPR, and 10 patients with psychogenic non-epileptic seizures (PNES). Diagnosis of GGE was established using International League Against Epilepsy (ILAE) criteria [31]. The diagnosis of PNES was confirmed with the consensus of at least two epilepsy specialists following video-EEG monitoring using criteria from a previous study [32]. Clinical data for all patients is summarised in Table 1, with a patient-specific breakdown in Supplementary Material Table S1. The use of this data for study was approved by the human research ethics committee of St Vincent’s Hospital Melbourne, where the data was obtained.

**Table 1:**
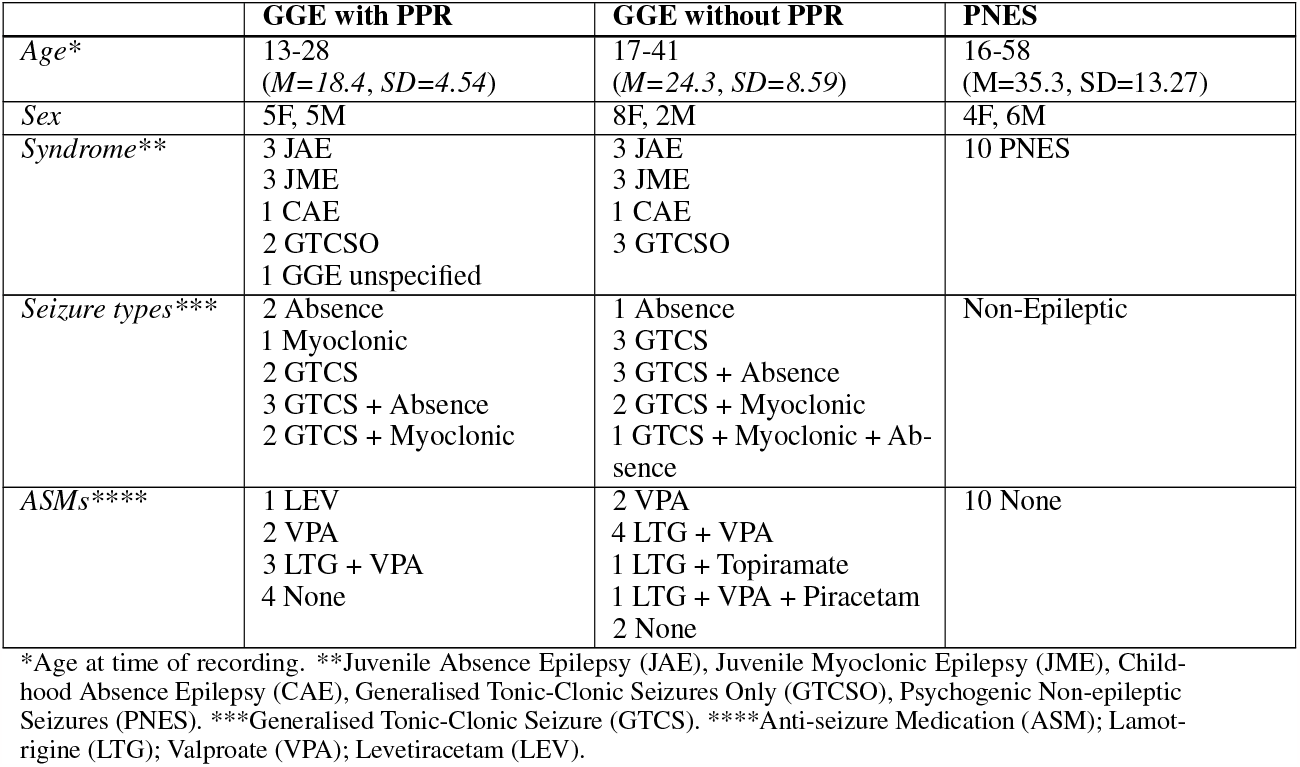
Summary of clinical data for participants in all groups.

#### 2.1.1 Healthy Controls

Healthy control EEG data was obtained from an open source repository, originally described and analysed in a study by Torkamani-Azar et al. [33]. Control data included ten (six female) healthy adults aged 22-45 (*M* = 30.25, *SD* = 6.95). For each control subject, we analysed 2.5 minutes of eyes-closed, resting-state EEG. The eyes-closed condition was chosen for consistency since our analysis focuses on stimulation epochs in which participants were instructed to close their eyes. More information regarding recording method is provided in the original study. Electrodes were selected to match our data, and sampling rate was conserved. The same EEG pre-processing and measure calculations were used for this control dataset.

### 2.2 Electroencephalography recording

Electroencephalography (EEG) data were acquired with a Compumedics Grael 4K-EEG system (Compumedics Ltd, Melbourne, Australia) or Siesta 802 in the case of ambulatory recordings. Scalp electrodes were placed according to the 10-20 international system. All EEGs were recorded with a bandwidth of 0.15-120Hz, with a sampling rate of 256Hz, including provocation techniques such as hyperventilation and intermittent photic stimulation (IPS) (as illustrated in Figure 1). Photo-paroxysmal response (PPR) was identified through visual inspection of the recordings, defined as epileptiform discharges triggered by IPS visible on the EEG. For clinical reasons, a number of participants, EEG recordings were also continued for a full day recording using the ambulatory recording system, sometimes capturing spontaneous seizures.

**Figure 1:**
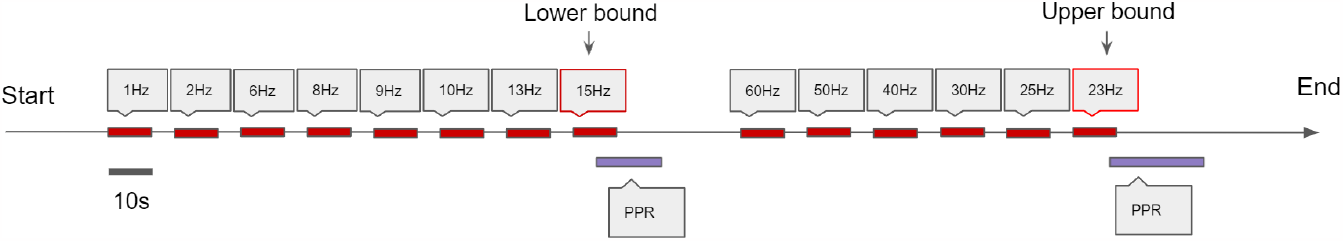
Schematic of the Photic Stimulation procedure. Stimulation epochs last for 10s with a 7s rest in between. The frequency of subsequent stimulations increase until a Photo-Paroxysmal Response (PPR) is triggered, and this activating frequency is designated the lower bound. Then, the technician starts from a high frequency and decreases the frequency of subsequent stimulations until another PPR is triggered, and this frequency is designated the upper bound. These lower and upper bounds are indicative of the range of frequencies that trigger a PPR response specific to that patient.

For GGE (PPR) participants, the ordering of stimulation frequencies often differed after their first PPR was observed; stimulation frequencies were sometimes repeated in an attempt to elicit PPR again, and sometimes alternated to high frequencies to determine an upper sensitivity range, as is convention [34]. It is not clear whether stimulation order effects exist, whether or not this depends on frequency, and whether refractory effects may exist after PPR that effects propensity for further PPR. Therefore, in an attempt to control for these confounding variables, we only analysed data before the first PPR. For GGE (no PPR) and PNES participants, stimulation frequency ordering was the same. In order to account for confounding effects of frequency-dependent responses, we only included stimulation epochs up to 15Hz in GGE (no PPR) and PNES groups in our analyses, as we observed PPR at or before 15Hz in all GGE (PPR) participants.

### 2.3 Pre-processing

Analyses were performed by importing raw data files from Profusion EEG 5 into MATLAB R2021a.

For variance and autocorrelation measures, we applied bandpass filters between 0.5 and 70Hz to reduce high frequency artifact and remove DC drift. For phase-based measures, this filtering step was ommitted, as more narrow bandpass filtering was later applied to calculate the measure.

To normalise the data, we focused on making the majority of the signal comparable between different channels with varying ranges, allowing tail values but without being overly skewed by the presence of artefacts distorting the re-normalised range. Therefore, we applied z-score normalisation using the native MATLAB `normalize` function, using the medianiqr argument to centre the data at 0 and ensure an inter-quartile range of 1, with a window length of two seconds. In GGE (PPR) participants, since only data before their first PPR was analysed, normalization was not affected by the high amplitude spiking during PPR.

### 2.4 Measures

To capture the expected rapid dynamics, measures generally used small windows with maximal overlap, and are described in detail below.

#### 2.4.1 Variance and autocorrelation function width (ACFW)

Variance and autocorrelation function width (ACFW) were calculated for each non-reference EEG channel (*n=19*). The variance of the signal was calculated for each channel with a moving window of two seconds (512 samples), with overlapping steps of 1 sample using the native MATLAB `movvar` function. To measure the autocorrelation function width (ACFW), using the same aforementioned window and overlap, the autocorrelation function was first calculated using MATLAB’s xcorr function, and the width in samples was measured at the half prominence of the function peak, centered at it’s maximum.

#### 2.4.2 Phase-based synchrony measures

To calculate phase-based synchrony measures, we obtain the phase-angle time series *φ*(*t*) (the instantaneous phase of each sample in the time-series) by constructing an analytic signal where the real component is the original time-series *x*(*t*), and the complex component is the Hilbert transform of the real signal ℋ[*x*(*t*)]. In other words, the analytic signal is the product of the envelope and instantaneous phase of the original timeseries,

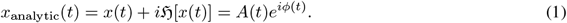

Note that this only holds if the original signal satisfies Bedrosian’s Product Thoerem, which requires that the signal is narrow-banded using a band-pass filter [35]. To do this, we filtered the normalised data using an elliptic filter, which is a zero-phase filter that circumvents phase shifting [36]. Specifically, a filter was constructed using MATLAB’s `designfilt` function, using the `bandpassiir` filter, with a filter order of 10, pass-band frequency limits according to the upper and lower limits of standard EEG frequency bands (delta: 1.5-4 Hz; theta: 4-8 Hz; alpha: 8-12 Hz; beta: 13-30 Hz; gamma: 30-70 Hz), a pass-band ripple of 3, and stop-band attenuation of 40.

#### Relative phase

Cosine of the Relative Phase (CRP: [37]) is calculated by taking the cosine of the instantaneous phase difference between two channels: *cos*(Δ*φ*(*t*)), where *φ*_*x*_(*t*) *− φ*_*y*_(*t*) obtained from the original channel timeseries *x*(*t*) and *y*(*t*) with the method previously explained for a given frequency band. This produces a measure ranging from *−*1 to 1, where magnitudes close to 1 denote high synchrony. Values close to *−*1 also arguably demonstrate a highly synchronous event, since this is produced when there is a phase difference of exactly *π*, known as an anti-phase relationship [37]. Measures were considered for each combination of non-reference channels (*n* = 19), resulting in *n*(*n −* 1)/2 = 171 unique combinations of channels.

#### Global and Posterior Synchrony

We also calculated the average synchrony between sets of channels using the average phase vector for a given sample. For *N* oscillators, with instantaneous phases *θ*_*j*_ for *j* = 1, 2, …*N*, their position can be represented on the unit circle by 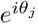. Then, the complex number *z* is the average of their positions,

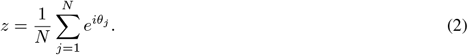

We found the magnitude of *z* using MATLAB’s native abs function; this produces a value from 0 to 1, where 0 represents no synchrony, or a uniform distribution, and 1 represents all phases being equal over all channels for a given time sample (see Fig. 2 for an illustration). We calculated the magnitude of the average phase vector over all 19 channels and designate this *global synchrony*, as well as over six posterior channels only (P3, P4, T5, T6, O1, O2) and designate this *posterior synchrony*.

**Figure 2:**
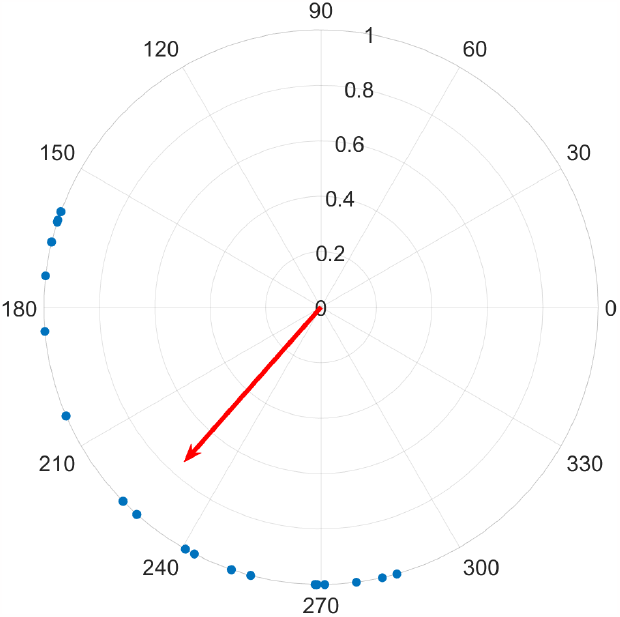
Each of 19 channels are shown on the unit circle as a blue dot, representing the instantaneous phase for a given time sample. The average phase is indicated by the direction of the red vector on the unit circle, and it’s length corresponds to their similarity, where a perfect alignment of phase would produce a length of 1, while a uniform distribution around the circle would be 0.

The magnitude of the mean phase vector can also be calculated over an epoch of time. For *N* oscillators over an epoch containing *S* samples, the instantaneous phases *θ*_*j,k*_ for channels *j* = 1, 2, …*N* and time samples *k* = 1, 2, …*S* can be represented on the unit circle by 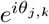. Then, the complex number 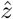 is the average of their positions,

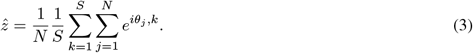

Again, the magnitude of this vector produces a value from 0 to 1, where 0 represents no synchrony, or a uniform distribution, and 1 results if all phases were equal between all channels over all time samples in the epoch.

### 2.5 Analysis strategies and statistics

#### 2.5.1 Clinical characteristics

Differences in ages between the four groups were evaluated using a one-way ANOVA (using the anova1 function from the MATLAB Statistics and Machine Learning Toolbox: SML). If significance was observed, follow-up multiple comparisons were evaluated using Tukey’s honestly significant difference (HSD) procedure (by passing the `stats` struct returned from the ANOVA into `multcompare` from the SML Toolbox).

Categorical clinical characteristics such as sex were examined between all four groups using Pearson’s chi-squared test of independence (using the `crosstab` function from SML). Differences in ASM, syndrome, and seizure type(s) were also evaluated using this test between the two GGE groups (with and without PPR) only, since PNES participants were not taking ASMs and were currently exhibiting non-epileptic seizures. Categories of ASM and seizure type(s) were considered distinct for each unique combination, as grouped in Table 1, since this allows for distinction of combinations that may be clinically significant.

The aforementioned tests were deemed significant with a *p*-value less than 0.05. Note that Tukey’s HSD procedure adjusts p-values for pair-wise multiple comparisons automatically.

#### 2.5.2 The photo-paroxysmal response

We compared time-frequency spectrograms of PPR events and spontaneous seizures for the four participants (P1, P2, P5, P6) for which both events were available. To generate the spectra, unfiltered epochs were fed to MATLAB’s `spectrogram` with window size of 256 samples, overlap of 128, and transform (nfft) length of 512 samples. Other PPR characteristics including PPR count, duration, and lower and upper frequency bounds are also measured for each participant.

#### 2.5.3 Measures

To investigate the hypothesis that photic stimulation perturbs brain dynamics towards a more excitable state that is measurable via changes to timeseries measures, we compared distributions of a number of measures during photic stimulation epochs compared to a control distribution. Substantial deviations from the control distribution suggests a shift in dynamics of the underlying system.

Photic stimulation distributions contained epochs before the first PPR in the GGE (PPR) group, or before the 15Hz stimulation in the GGE (no PPR) and PNES groups. We also examined an additional pre-PPR epoch distribution in the GGE (PPR) group only, containing the closest full stimulation epoch before (ie not overlapping) the first PPR event.

Since sufficient quality resting state data from the photic stimulation recordings were not available, we compared our data to distributions of healthy controls during eyes-closed, resting EEG. To create the control distributions, we randomly sampled 10s epochs from the full measure timeseries, with equal representation from the individuals of the cohort. This constructed distribution emulated the process of pooling 10s stimulation epochs in the epileptic cohort groups.

Differences between pairs of group distributions were evaluated using permutation-based hypothesis tests, which are non-parametric tests that do not make assumptions about the distribution of the data [38]. This process involves creating a joint distribution of two considered groups, and performing the test statistic of interest on random permutations of the joint distribution (where each permutation is equally as likely as another) to construct a distribution of test statistics (*n* = 10, 000) that represents the null result distribution. The test statistic between the original separate distributions is then compared to the null distribution, where the *p*-value is the fraction of permutation tests that are at least as extreme as that test statistic. A pseudocount (addition of one to the fraction numerator) is included to avoid *p*-values of zero (where the test statistic is never surpassed by permutation values). This is possible because we used a subset of the possible permutation set (though subsets of sufficient size have been shown to be accurate) [39]. In our case, this single test statistic value was the average value of a test distribution constructed from permutation subsets of the two considered distributions. From the central-limit theorem, this test statistic distribution will be normal with the mean at its centre. We created confidence intervals around this statistic by extracting the 2.5th and 97.5th percentile of the test distribution.

We used the Kolmogorov-Smirnov test statistic for continuous measures, which evaluates the difference between empirical cumulative distribution functions (CDF), using rank-based comparisons. This is appropriate considering variance and synchrony measures demonstrated non-normal distributions through inspection of q-q plots. The Kolmogorov-Smirnov test identifies small differences in large datasets as significant even when differences seem trivial [40]; therefore, permutation samples of size 500 were taken, in accordance with sample size recommendations [41]. Chi-squared tests were used as the test statistic for auto-correlation function width, which is a discrete measure.

We also investigated whether measures of synchrony would change with photic stimulation. The relative phase measure contained significant degrees of freedom (unique channel pairs = 171, frequency bands = 5, resulting in 855 potential comparisons), so we investigated whether factor analysis could reduce the parameter space by identifying channels that demonstrated the greatest variation. Factor analyses of the GGE (PPR) stimulation epochs (using factoran from SML, with 5 component factors to generate the maximum likelihood estimate) were carried out. This analysis was performed for each frequency band separately to investigate whether variation was also frequency-band dependent.

To further investigate whether measures varied by channel, and whether this is apparent in posterior channels reflecting underlying posterior pathophysiology, we compared posterior and global synchrony distributions to the corresponding control distributions. Differences were evaluated using the same permutation based hypothesis tests described above, using the two-sample Kolmogorov-Smirnov test as the test statistic, since synchrony values are continuous.

Bonferroni correction, considered the most conservative multiple-comparison correction, was carried out (the standard *p*-value was divided by 56: a total of 12 comparisons for variance and autocorrelation distributions, and 5 frequency bands x 4 distributions x 2 sets of channel groups in the synchrony analyses), requiring a *p*-value less than 0.00096 for significance.

## 3 Results

### 3.1 Participant clinical characteristics

One-way ANOVA identified a difference in age between at least two groups (*F* (3, 36) = 6.77, *p <* 0.001). Posthoc analyses revealed that the following significant differences existed: the GGE (PPR) group had lower ages compared to the PNES group (*p <* 0.001, 95% *C*.*I*. = [*−*27.77, *−*6.03]) and compared to controls (*p* = 0.018, 95% *C*.*I*. = [*−*23.42, *−*1.68]). The GGE (no PPR) group also had lower ages compared to the PNES group (*p <* 0.046, 95% *C*.*I*. = [*−*21.87, *−*0.13]). We did not reject the null hypothesis that group and sex are independent (*χ*^2^(3) = 3.58, *p* = 0.31); therefore, groups did not differ in sex ratio.

We also investigated whether ASM, syndrome, and seizure type differed between the two GGE (with and without PPR) groups. We did not reject the null hypothesis of independence between group and ASM (*χ*^2^(5) = 3.81, *p* = 0.58), syndrome (*χ*^2^(4) = 3.53, *p* = 0.47), or seizure type (*χ*^2^(5) = 1.87, *p* = 0.87); therefore, these groups did not differ in the spread in these variables (see Table 1 for a breakdown of categories).

### 3.2 The photo-paroxysmal response (PPR)

PPR featured spike-wave morphology in all channels and lasted between 0.6-28 seconds (see Figure 3 for an example). As can be seen in Table 2, GGE patients with PPR exhibited individualised responses to photic stimulation, including differences in their photosensitivity ranges as well as PPR length.

**Table 2:** Participant-specific information about PPR events.

| Participant | Lower and upper frequency (Hz) bounds | PPR count (n) | Average PPR length (s) |
| --- | --- | --- | --- |
| 1 | 15-23 | 2 | 22.69 |
| 2 | 15-20 | 2 | 2.93 |
| 3 | 15-23 | 2 | 1.58 |
| 4 | 18-30 | 2 | 0.81 |
| 5 | 8-30 | 2 | 4.00 |
| 6 | 8-25 | 2 | 2.25 |
| 7 | 13 | 1 | 1.47 |
| 8 | 9-30 | 5 | 3.26 |
| 9 | 13-50 | 3 | 2.15 |
| 10 | 13-25 | 5 | 1.55 |

**Figure 3:**
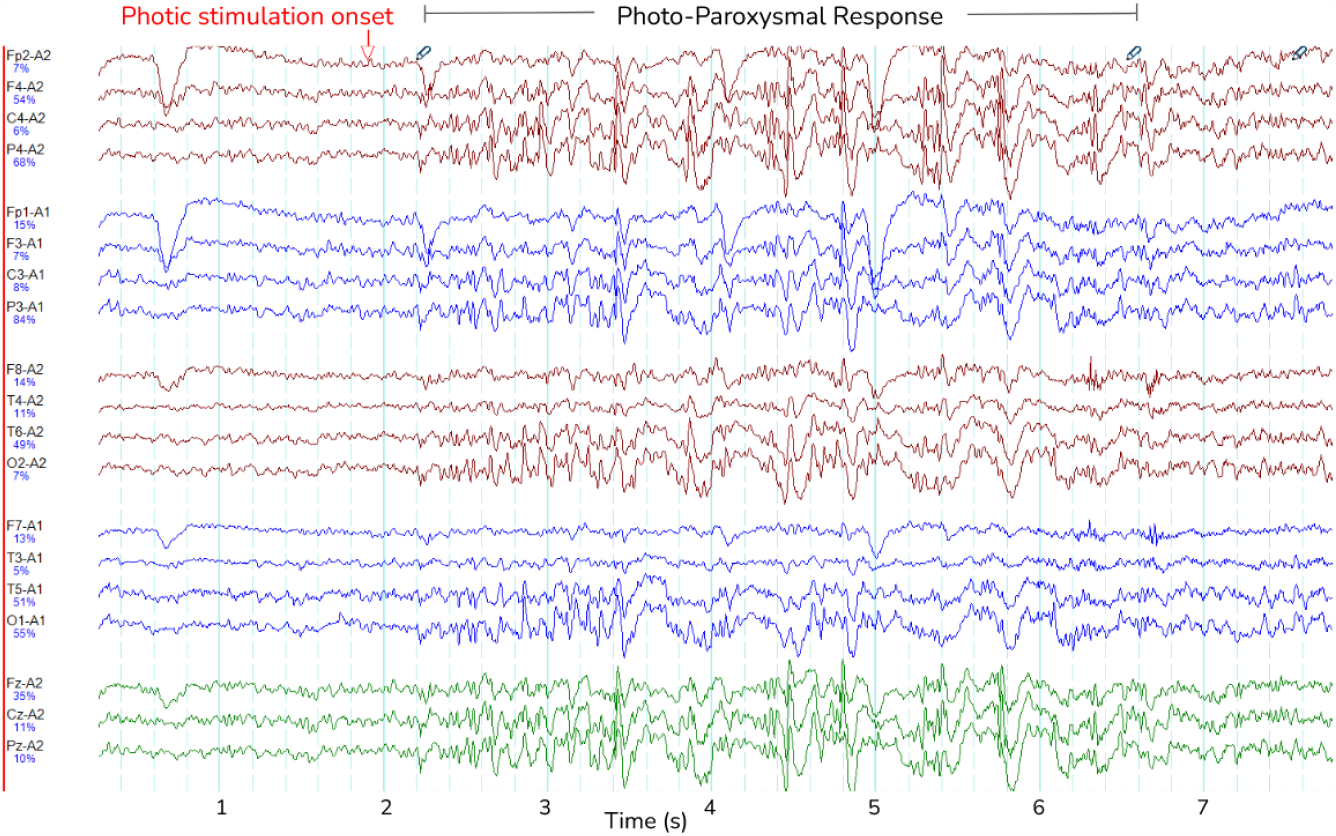
Photo-paroxysmal Response (PPR). The figure shows a typical PPR lasting approximately 5 seconds, visible as spike-wave morphology. Lines are different channels as indicated on the y-axis, and time is indicated in seconds on the x-axis.

In four participants, spontaneous seizures were observed hours after photic stimulation. We compared the transition to PPR events and spontaneous seizures by their time-varying frequency spectra by visual inspection (Fig. **??**). Though spectra varied according to the individual, we observed changes to the distributions of power per frequency bin at the transition, including increased power especially in delta and theta frequencies. The spectral dynamics of PPR and seizure transitions appear similar.

We inspected spectrograms over participants’ stimulation trains and often observed increased power in the base frequency of stimulation, as well as its harmonics (see Supp. Fig. 2). We observed this in all participant groups during photic stimulation. Comparison is difficult due to varying data quality and different stimulation frequency ordering in the photosensitive group. The visibility of these bursts of increased power varied substantially by participant, though we report that it was most visible in the occipital channels. Interestingly, they are found to exist even during PPR (see Supp. Fig. S2).

### 3.3 Measures

We examined whether measures could track individualised responses to perturbations by examining the timeseries of measure values from GGE (PPR) participants as a PPR approaches.

#### 3.3.1 Variance and autocorrelation

From inspecting individual stimulation trials, we observed an increase in variance during stimulation epochs close to a PPR (see Fig. 4 and Supp. Fig. S4 for further examples).

**Figure 4:**
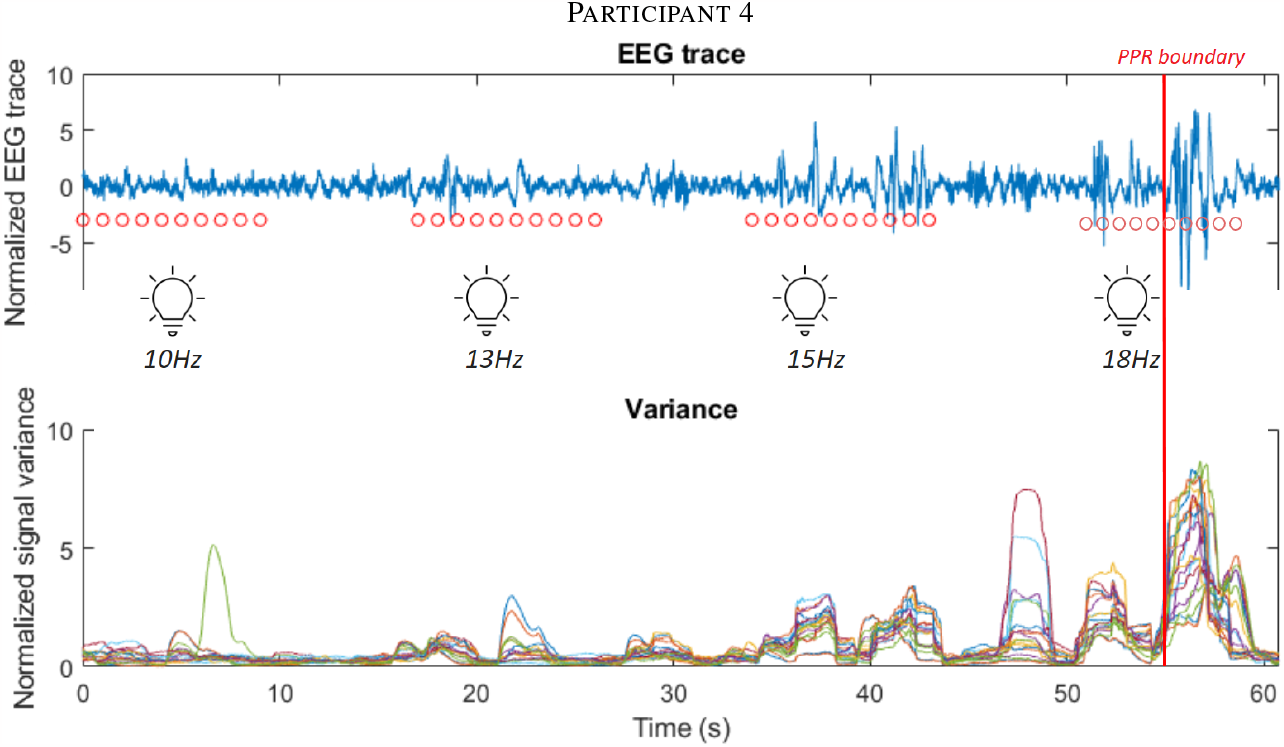
The first subplot shows the EEG trace (blue line) of participant 4 in the leadup to a PPR, visible as high amplitude spiking activity after the marked PPR onset boundary (vertical red line). Consecutive red dots mark 10 seconds of photic stimulation for the frequency (labelled below them). Below, the corresponding variance of the signal is displayed, using the shared time axis (y-axis), where different electrode channels are displayed as different coloured lines. Observe the increasing variance close to the boundary. See Supp. Fig. S4 for further participant examples.

We analyse whether these measures may track cortical excitability using group-wise comparisons. To do this, we compared the distributions of variance and autocorrelation function widths (ACFW) during photic stimulation epochs between groups. Values were pooled over all 19 electrode channels and the 10 participants in each of the four groups. We compared the cumulative distribution functions of these measures to resting healthy controls, and observed a greater relative proportion of high values in all groups during photic stimulation (see Fig. 5). Significant differences in empirical cumulative distribution functions (CDF) was found compared to controls in all groups during stimulation, as well as in the the GGE (PPR) group just prior to their first PPR (‘pre-PPR’), where cortical excitability was hypothesised to be greatest (see Table 3 for a breakdown of statistical test outputs). There was no significant differences between the photosensitive GGE (PPR) group and the nonphotosenstive epilepsy groups.

**Table 3:** Statistical results.

| Distribution 1 | Distribution 2 | Variance | Autocorrelation Function Width |
| --- | --- | --- | --- |
| GGE (PPR) stim | GGE (PPR) pre-ppr | $D = 0.19^*$<br>$95\%CI = [0.14, 0.25]$ | $\chi^2 = 44.86^*$<br>$95\%CI = [27.00, 66.33]$ |
| GGE (PPR) stim | GGE (no PPR) stim | $NS$ | $NS$ |
| GGE (PPR) stim | PNES stim | $NS$ | $NS$ |
| Controls resting | GGE (PPR) stim | $D = 0.16^*$<br>$95\%CI = [0.12, 0.20]$ | $\chi^2 = 368.42$<br>$95\%CI = [319.78, 419.03]$ |
| Controls resting | GGE (no PPR) stim | $D = 0.15^*$<br>$95\%CI = [0.11, 0.19]$ | $\chi^2 = 226.86$<br>$95\%CI = [185.66, 269.26]$ |
| Controls resting | PNES stim | $D = 0.12^*$<br>$95\%CI = [0.08, 0.15]$ | $\chi^2 = 272.92$<br>$95\%CI = [227.15, 319.27]$ |

Significance is evaluated through permutation ( $n = 10,000$ ) hypothesis tests of the test statistic where '\*' denotes significance (adjusted for multiple comparisons using bonferroni correction). $NS$ denotes no significant difference. $D$ is the Kolmogorov-Smirnov test statistic. $\chi^2$ is the Chi-squared test statistic. $95\%CI$ obtained through the 2.5th and 97.5th percentiles of the test statistic distribution constructed through permutation re-sampling ( $n = 10,000$ ).

**Figure 5:**
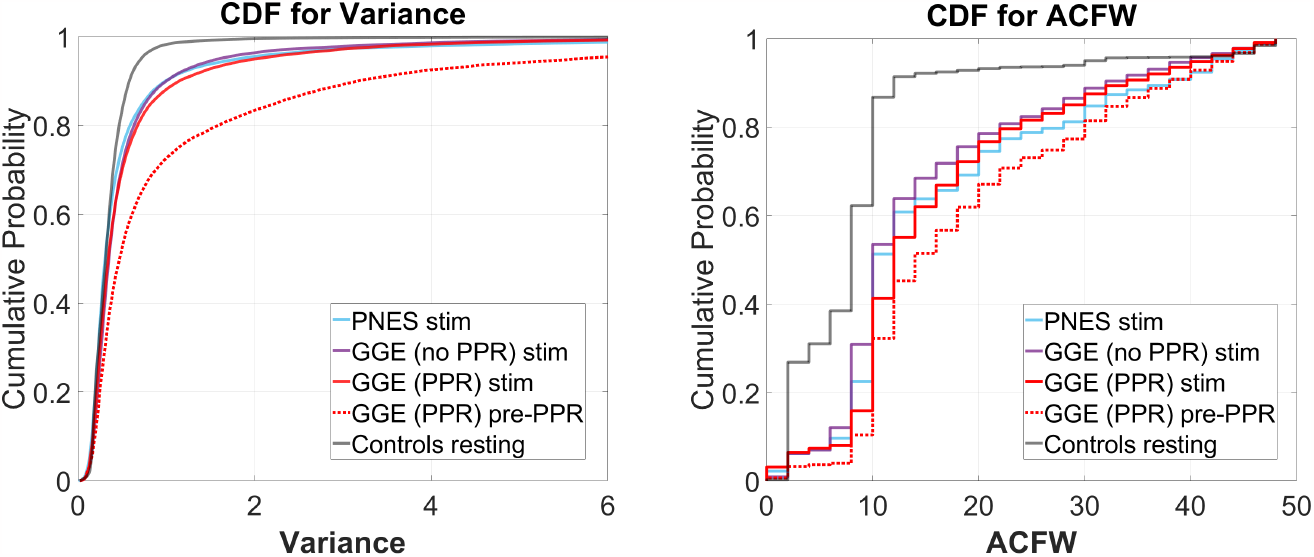
**(A)** variance (left plots) and **(B)** autocorrelation function width (ACFW; right plots), where the x-axis is the values for each measure, and the y-axis is the probability of observing a corresponding x-value and all values less than it. Each coloured line represents a different group’s distribution, pooled over all participants and channels for selected epochs (refer to legend). Right-shifted lines indicate distributions with larger proportions of high values. Observe how all groups show right-shifts compared to the control (grey) line. Also observe how, for the GGE (red) group, their distributions shift further to the right, in the epoch just before PPR (dotted line labelled as ‘pre-PPR’), compared to earlier stimulation epochs (solid line labelled as ‘GGE (PPR) stim’).

#### 3.3.2 Synchrony Relative Phase

To investigate whether changes in synchrony may indicate increased cortical excitability, we constructed phase diagrams of relative phase values in stimulation epochs leading to a PPR. In some cases, we observed greater clustering in epochs close to a PPR, indicating that a particular channel pair was becoming more synchronous (marked also by longer mean vector lengths - see Fig. 6). However, clustering magnitudes varied considerably between individuals and depending on combinations of factors like channel pair, frequency band, and time to PPR, suggesting considerable individual variation. Magnitudes tended to correlate positively between frequency bands, though lower frequency bands (delta, theta, alpha) tended to have ranges extending to higher values, compared to beta and gamma which tended to have lower values overall, likely related to different degrees of freedom from bandwidth ranges).

**Figure 6:**
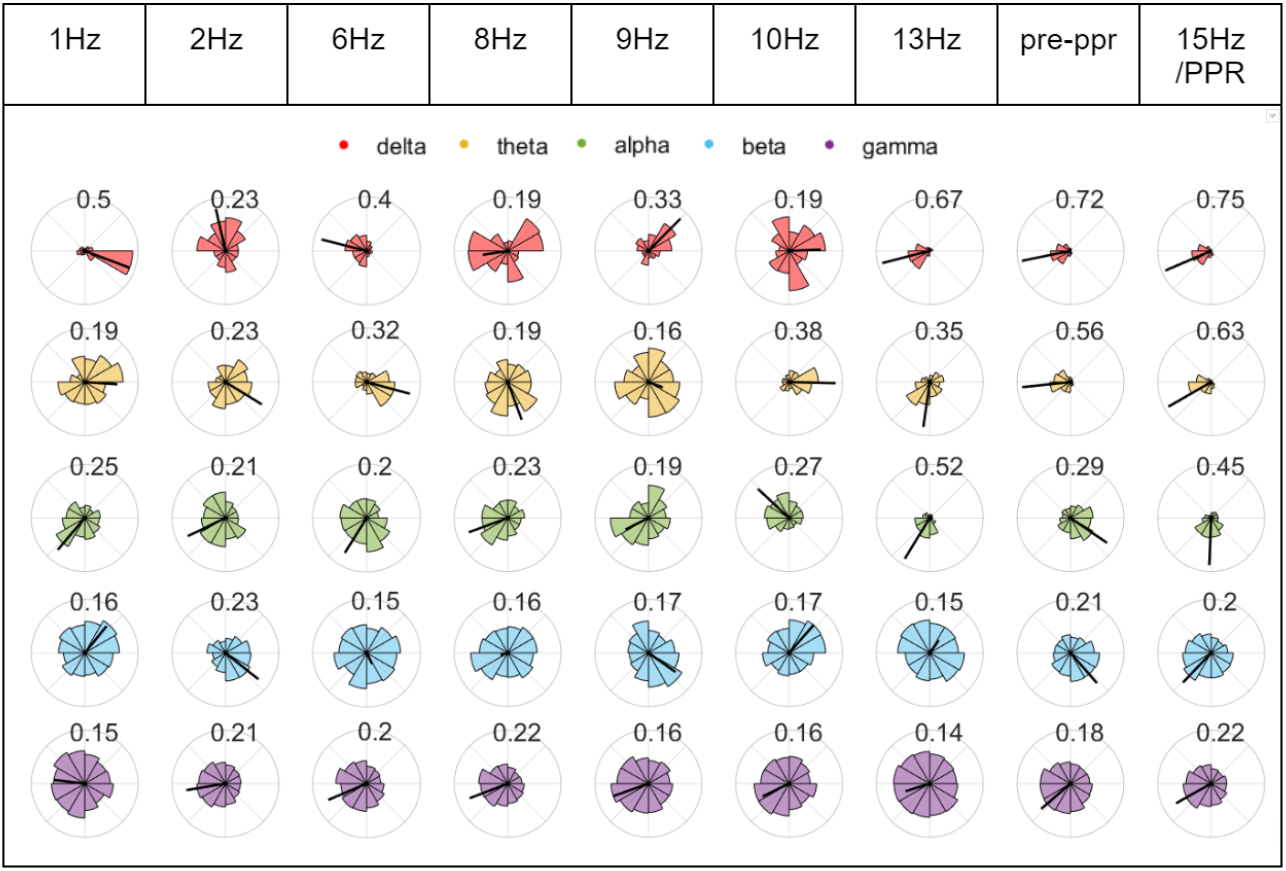
Chronological phase histograms during P1’s stimulation train. Each plot represents 10s of binned relative phase difference values between O1 and Fz for epochs leading to the first PPR. Each bin has a direction from 0 to 2pi around the unit circle, and a length indicating the relative frequency of values, ranging from 0 to 1. The radius of each circle plot varies to assist visualisation, and the the mean relative phase amplitude (time-varying synchrony) is indicated by a black line whose length is labelled above each plot and also ranges from 0 to 1, where 1 indicates greater clustering of the values in the plot. Each row corresponds to relative phase values within different frequency bands (see legend).

To investigate whether certain channels may demonstrate greater change in synchrony, a factor analysis was performed on relative phase values during stimulation from the GGE (PPR) participants. In this analysis, the data from each EEG channel are assumed to depend on a linear combination of latent components, which attempt to capture the variance of the data along their axes, (similar to Principal Component Analysis). Each channel’s factor loadings for each component axis indicate the amount of variance in the data that was captured along that component. The three dominant component factors separated a subset of posterior channels, which demonstrated the greatest variance along these factor axes (see Fig. 7). This result was relatively consistent over all frequency bands (see Supp. Fig. S5).

**Figure 7:**
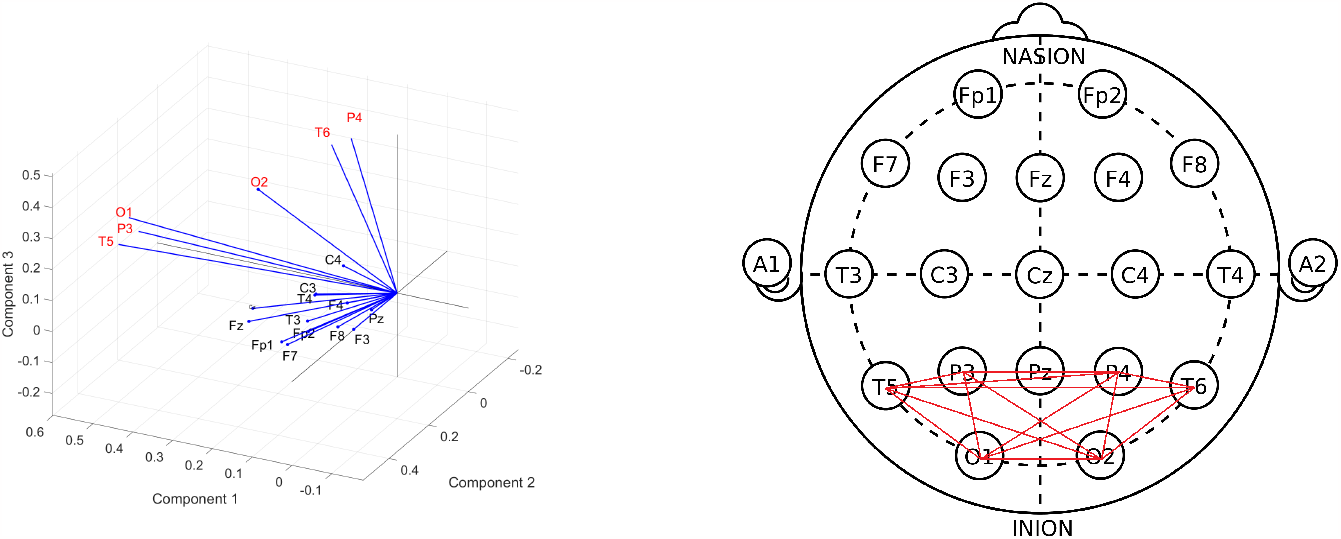
**(A)** Biplot of factor loadings (blue vectors) on the three dominant components (axes), generated from a factor analysis of cosine of the relative phase values during photic stimulation, pooled over GGE (PPR) participants. Each channel’s factor loadings for each component axis indicate the amount of variance in the data that was captured along that component. Channels with the highest factor loadings, or in other words demonstrated the greatest variance, along the component axes are P4, T6, O2, O1, P3, T5 (labelled in red text). This suggests that the greatest change in the data was demonstrated by this subset of channels. Similar results were obtained when repeated in other frequency bands (see Supp. Fig. S5). **(B)** Subgraph of posterior channels.

#### Global and Posterior Synchrony

To investigate changes in synchrony in broader brain regions, we compared global and posterior synchrony in groups during photic stimulation compared to controls. Significant differences in empirical cumulative distribution functions (CDF) was found compared to controls in some frequency bands and groups for both global and posterior synchrony (see Fig. 8 for an illustration of the CDFs, and Table 4 for a breakdown of statistical test outputs). For global synchrony, stimulation groups differed significantly to controls in some frequency bands, including gamma in all groups, with an overall reduced probability of high coherence values (see Fig. 8). In comparison, posterior distributions showed increased probability of high coherence values in stimulation groups compared to controls, reaching significance for all frequency bands except the GGE (PPR) stimulation gamma distribution. There was no significant differences between the pre-PPR distribution and stimulation distribution for GGE (PPR) group for global or posterior synchrony.

**Table 4:**
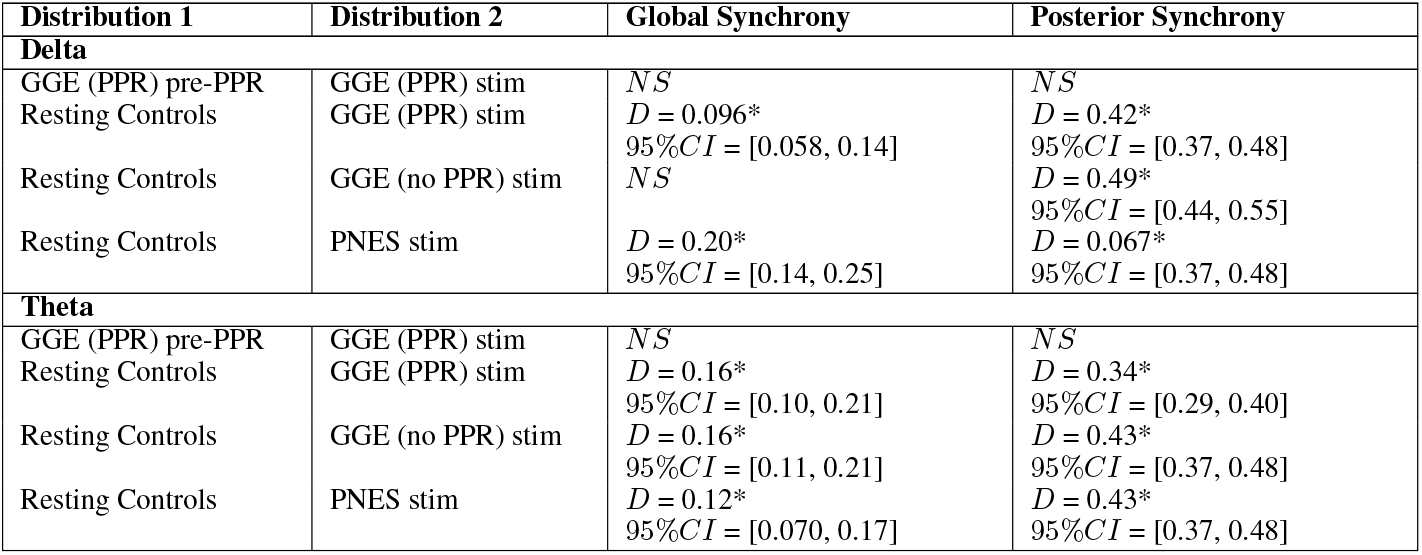
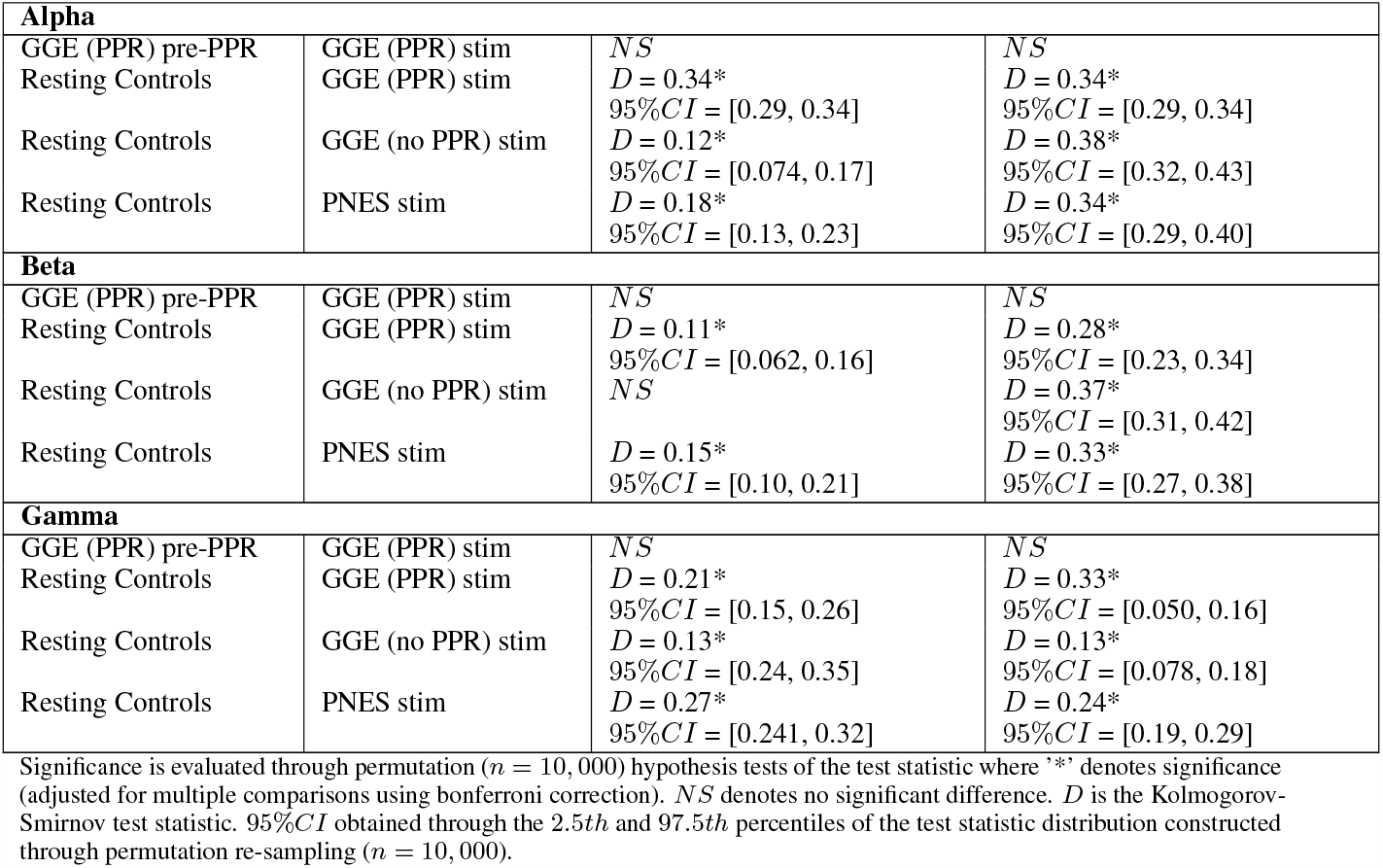
Statistical results.

**Figure 8:**
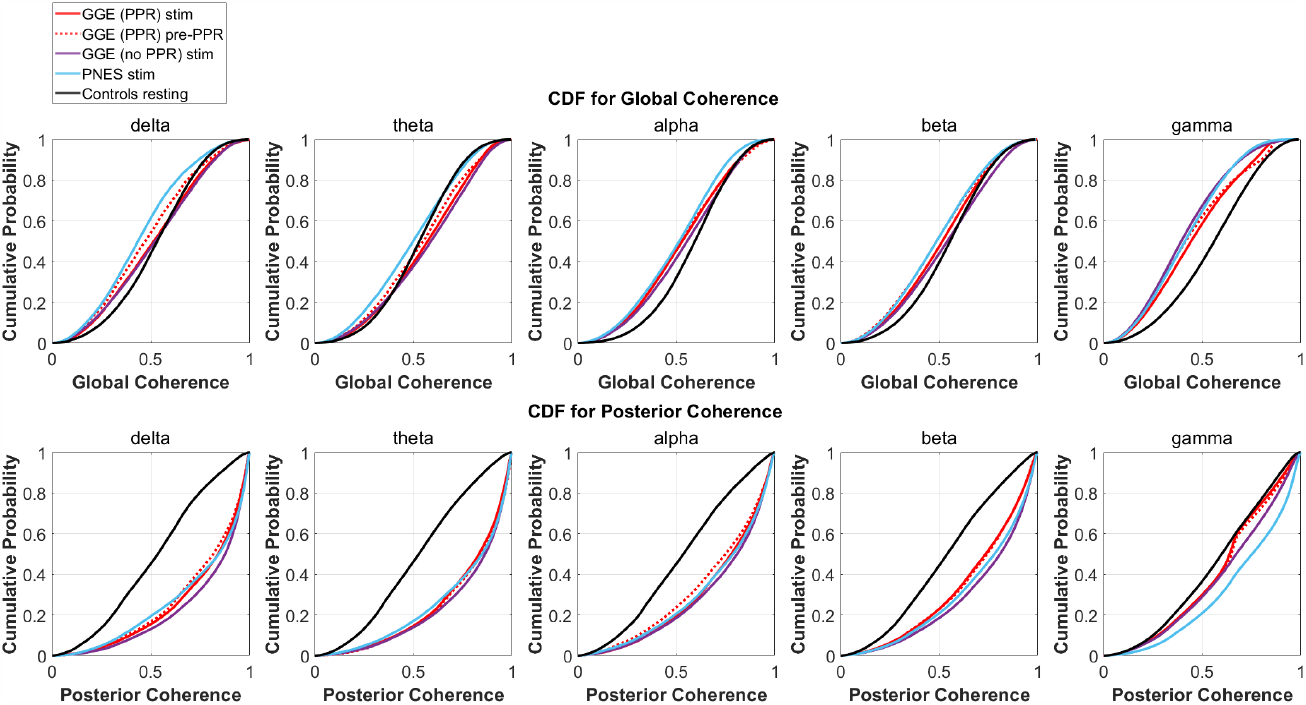
Empirical cumulative distribution functions (CDF) for coherence in different groups (different coloured lines, see legend), within different frequency bands (columns). The x-axis is coherence calculated as the mean phase vector magnitude over a set of channels. The y-axis is the cumulative probability for the corresponding x-values. From visual inspection, there is evidence for decreased global synchrony during stimulation in the first row where left-shifts are observed compared to controls (black line), most evident in the gamma band. Increased posterior synchrony is evident in the second row in all frequency bands, with right shifted CDFs for groups during stimulation (coloured lines) compared to controls (black line). See Table 4 for a breakdown of significant differences. **(A)** CDFs for Global Coherence (mean magnitude over all channels). **(B)** CDFs for Posterior Coherence (mean magnitude over posterior channels only).

Distinction between each group’s stimulation distribution is not seperable from inspection of CDF distributions, though the presence of differences relative to controls across stimulation groups suggests a group-independent effect of the photic stimulation on relative global and posterior synchrony, that is not specific to a particular group.

## 4 Discussion

### 4.1 Summary

The primary finding of this study was that the response to perturbation by photic stimulation was observable via changes in EEG measures. Specifically, distributions of variance and autocorrelation function width (ACFW) shifted to higher values just before photo-paroxysmal response (PPR), indicating greater cortical excitability close to seizure transitions, as hypothesised. In all epilepsy groups, phase coherence distributions differed to controls. As expected, posterior channels demonstrated the greatest variation in synchrony. Greater changes were sometimes observed in epochs close to PPR for some individuals, but there was no significant difference at a group level. This is not surprising considering the patient-specific nature of responses, and we suggest future research apply individualised analyses utilising repeated-trials. We have demonstrated measurable changes of excitability across different groups of epilepsy patients using photic stimulation, particularly photosensitive patients in the lead up to seizure transitions consistent with previous literature [12; 11; 7]. Therefore, we argue that PPR is a valid proxy for studying seizure transitions. This human model is a safer alternative to inducing seizures, and can be reliably induced via photic stimulation. We argue this model has great potential for developing new diagnostic methods and evaluating treatments for epilepsy.

### 4.2 Relevance and significance of findings

At a group level, we observed greater proportions of higher variance and autocorrelation distributions in all group distributions during photic stimulation compared to resting controls (see Fig. 5). Individually, patients demonstrated considerable variation in response values, particularly in the photosensitive group, motivating our group-level analysis to understand overall trends. Values tended to be highest in the photosensitive group, compared to the non-photosensitive generalised epilepsy group, and patients with psychogenic non-epileptic seizures displayed the lowest values; however, this difference was not found to be significant in the group-level analysis. As expected, the photosensitive group demonstrated further increases just before photo-paroxysmal response.

We interpret the shift in distributions as a data-driven illustration of underlying dynamical regime changes close to seizure transitions. These observations are consistent with predictions of increased system summary statistics before state transitions in dynamical systems, together with increased responses to perturbation [12]. These early warning signs of transitions have been observed before seizure transitions [7; 42]. Though, our data differs in that it reflects changes to the actively evoked perturbation response, instead of just passive observations of change. Studies have elicited active perturbation using invasive EEG for focal epilepsy and observed similar warning signs [11; 43], while we report it here using non-invasive perturbation methods in generalised epilepsy. We argue that the lead-up to PPR demonstrates similar changes in dynamics to seizure transitions, making it a good proxy for studying seizure transitions. Further, PPR demonstrates characteristic spike-wave morphology, and similar spectral content to spontaneous seizures visible in spectrograms. Increased power was evident in a broad range of frequencies including the 2-5Hz activity characteristic of seizure activity [34].

We speculate that the increased variance before PPR, is associated with an increased firing activity of neurons in synchrony, considered to underpin seizure activity. Increased autocorrelation width suggests longer lasting increases to self-similarity [12], changes to the periodicity in the signal, decreased dimensionality [44], and possibly increased variance. We also understand that for cyclo-stationary signals, the autocorrelation function is equal to the absolute value of the square of its Fourier transform, in which case increased width of the major peak may reflect changes to the ratio of power in various frequency ranges. We observed distinct changes in power at the frequency of stimulation and their harmonics, which supports this explanation (see Supp. Fig. 2). It could also reflect a breakdown in the signals’ stationarity, which is to be expected during a change in dynamical regime. In sum, it is difficult to disentangle the exact mechanism behind these changes, although they do reflect changes in brain dynamics.

Compared to resting controls, all groups demonstrated significantly reduced gamma coherence when averaged over all channels (see Fig. 8) but increased coherence when averaged over only the posterior subgroup in nearly all frequency bands (see Fig. 7. We also found the majority of change in phase clustering (cosine of the relative phase) values occurred in the posterior subgroup via a factor analysis. This evidence of increased posterior synchrony is well established in the literature [6; 24; 45; 23], including in healthy individuals [46], suggesting recruitment of the visual system that is arguably enhanced to flicker in particular. It is likely that subsequent aberrant propagation of posterior synchronisation due to abnormal gain control results in PPR [23]. However, we were not able to find substantial group differences between photosensitive and non-photosensitive distributions during stimulation, or between the photosensitive group prior to PPR and earlier epochs, which suggests the measure was not able to indicate increased cortical excitability, as was found in ECoG data [11]. We did, however, observe substantial individual variation, and multiple examples of increased coherence close to PPR in some individuals, channels, and frequency bands (see Fig. 6). Therefore, we encourage future work to conduct individualised, frequency-specific evaluations of multiple stimulation trials to elucidate enhanced responses in photosensitive individuals.

### 4.3 Limitations

In this retrospective study, there were several limitations. Because we did not have appropriate resting state recordings, we compared data to resting controls. However, ideally, each individual would be compared to their own resting or baseline distribution to ascertain the true response to perturbation. Another issue was the considerable within-group variability that we observed, suggesting that the population distribution for each epilepsy group was itself heterogeneous and would overlap each other considerably. We observed frequency-dependent differences in the response to perturbation, consistent with previous research that generally finds marked responses in the 10-20Hz range [34], and in harmonic frequencies of the PPR-evoking frequency (activation frequency) [24]. However, it would not have been appropriate to perform a group-level frequency-dependent analysis, because of confounders like ordering effects due to variation in stimulation frequency order (which could be mitigated through randomisation), possible refractory effects from PPR events, and individualised responses to perturbation.

### 4.4 Future work

The applications for photic stimulation are three-fold: 1) it can be used to diagnose photosensitive epilepsy; 2) it can be used as a therapeutic tool for epilepsy patients to monitor treatment efficacy; and 3) it can be used as a perturbation paradigm for studying brain dynamics.

Photic stimulation may continue to be used for its current diagnostic purposes for epilepsy. However, we recommend for research purposes, that multiple trials within a single session may be necessary to obtain sensitive results. Multiple trials will also enable the construction of robust, patient-specific profiles of frequency-dependent responses to perturbations.

These profiles would be clinically significant in that changes to the initial measurements could be used to evaluate the efficacy of medications, which are understood to alter cortical excitability as their mode of action [47]. They could also be used to evaluate the safety of medication withdrawal, and create standardised clinical guidelines, by indicating whether drug removal has shifted patients to regimes of higher cortical excitability and, therefore, seizure risk [11; 43], and monitor disease course in the long-term. Also, it should be noted that observing measurable responses to perturbation from photic stimulation is not limited to photosensitive individuals. We observed responses to perturbation in all groups, regardless of epilepsy diagnosis (GGE or PNES), syndrome, seizure type, age, sex, or ASM. We conclude that the response to perturbation may be observable in the general epilepsy population, suggesting that the photic stimulation paradigm offers a great potential framework for studying seizure dynamics at large.

We wish to emphasise the potential for photic stimulation to be used as a perturbation paradigm for studying general brain dynamics. Previous, studies have already shown that greater responses to perturbation are associated with elevated cortical excitability relating to seizure risk in focal epilepsy [7], with cohorts of patients undergoing pre-surgical evaluations [17; 13] and with severe treatment-resistant epilepsy [16; 19]. Like the implanted electrical stimulation in these studies, photic stimulation enacts a perturbation. Compared to these surgical methods, it can be considered a much safer method for studying the transition to seizure, as well as compared to non-surgical alternatives like seizure provocation during hospitalisation via medication withdrawal and sleep deprivation. In fact, this non-invasive method could potentially be used at home, only requiring an EEG headset, appropriate software, and a monitor for administering the appropriate photic stimulation. Changes to an individual’s baseline at any frequency could indicate meaningful differences to cortical excitability and seizure risk, regardless of the type of epilepsy. Also, as the response to photic stimulation differs in other diseases including migraine [48], schizophrenia [49], alzheimer’s disease and mild cognitive impairment [50; 23], its relevance as a research paradigm to studying wider diseases of brain structure or function is significant, in particular cortico-thalamic circuitry dysregulation in which many brain diseases are implicated [51; 52]. Furthermore, photic stimulation could be used to observe changes in cortical excitability in healthy individuals under initiating factors [34; 23] like alcohol withdrawal and sleep deprivation, to provide further insight to acute symptomatic seizures of people without epilepsy and human brain dynamics in general.

### 4.5 Conclusion

We found that the response to photic stimulation was a valid non-invasive perturbation to brain dynamics. This paper represents photic stimulation data from photosensitive generalised epilepsy patients, patients with GGE who are not photosensitive, and patients with psychogenic non-epileptic seizures, compared to resting controls. At a group level, consistent with the hypothesis that PPR induces a change in brain state, we observed clear changes in group signal statistics including the CDF of the variance and autocorrelation. We further investigated changes in synchrony measures, and could not find consistent group-level patterns of change that correlated with the transition. However, we did find posterior increases in synchrony consistent with past research. coheren that decreased global coherence together with increased posterior coherence may relate to the underlying pathophysiology of the PPR. In conclusion, EEG time-series analysis was able to capture changes in cortical excitability during active perturbation, via biomarkers that track brain dynamics before state transitions. Photic stimulation demonstrates a non-invasive human method for studying seizure transitions, and potentially monitoring and evaluating treatments in epilepsy and other brain diseases.

## Notes

### Competing Interest Statement

The authors have declared no competing interest.

### Summary of Updates

All sections including the abstract, introduction, methods and results sections were revised to clarify statements, fix typos, and at times restructured for clarity. All figures were re-uploaded with higher resolution and sometimes revision of content for clarity. Figure 3 was added. Methods section was updated with explanations of permutation resampling significance testing, and the corresponding results tables (3 and 4) were also updated accordingly. Supplementary material was also revised including inclusion of higher resolution images and clarification of existing figure captions and addition of supplementary figure 2.

