## Supplementary Material for "Response to photic stimulation as a measure of cortical excitability in epilepsy patients"

### 5 Supplementary material

Table S1: Breakdown of clinical data for individuals by diagnosis

| Participant | Age* | Sex | Syndrome** | Seizure types*** | ASM**** |
|---|---|---|---|---|---|
| *Participants with genetic generalised epilepsy and photo-paroxysmal response* | | | | | |
| 1 | 15 | F | JAE | GTCS, Absence | LEV |
| 2 | 16 | F | JME | GTCS, Myoclonic, Absence | None |
| 3 | 17 | F | CAE | Absence | LTG, VPA |
| 4 | 18 | M | JAE | Absence | VPA |
| 5 | 26 | F | JAE | GTCS, Absence | LTG, VPA |
| 6 | 17 | M | GTCSO | GTCS | VPA |
| 7 | 28 | M | CAE | GTCS, Absence | None |
| 8 | 13 | F | GTCSO | GTCS | LTG, VPA |
| 9 | 16 | M | JME | GTCS, Myoclonic | None |
| 10 | 18 | M | GGE unspecified | Myoclonic | None |
| *Participants with genetic generalised epilepsy, without photo-paroxysmal response* | | | | | |
| 11 | 25 | F | JME | GTCS, Myoclonic | LTG, VPA |
| 12 | 23 | F | JAE | Absence | LTG, Piracetam, VPA |
| 13 | 23 | M | GTCSO | GTCS | None |
| 14 | 21 | F | JME | GTCS, Myoclonic, Absence | LTG, VPA |
| 15 | 15 | F | JAE | GTCS, Absence | None |
| 16 | 16 | F | JAE | GTCS, Absence | VPA |
| 17 | 37 | F | GTCSO | GTCS | LTG, VPA |
| 18 | 25 | M | CAE | GTCS, Absence | LTG, VPA |
| 19 | 41 | F | JME | GTCS, Myoclonic | LTG, Topiramate |
| 20 | 17 | F | GTCSO | GTCS | VPA |
| *Participants with psychogenic non-epileptic seizures* | | | | | |
| 21 | 30 | F | PNES | | None |
| 22 | 30 | M | PNES | | None |
| 23 | 58 | F | PNES | | None |
| 24 | 31 | F | PNES | | None |
| 25 | 40 | F | PNES | | None |
| 26 | 30 | M | PNES | | None |
| 27 | 30 | M | PNES | | None |
| 28 | 16 | M | PNES | | None |
| 29 | 30 | M | PNES | | None |
| 30 | 58 | M | PNES | | None |

* Age at time of recording ** Juvenile Absence Epilepsy (JAE), Juvenile Myoclonic Epilepsy (JME), Childhood Absence Epilepsy (CAE), Generalised Tonic-Clonic Seizures Only (GTCSO), Psychogenic Non-epileptic Seizures (PNES) *** Generalised Tonic-Clonic Seizure (GTCS) **** Anti-seizure Medication (ASM); Lamotrigine (LTG); Valproate (VPA); Levetiracetam (LEV)

S2A: PARTICIPANT 1 (GGE PPR)

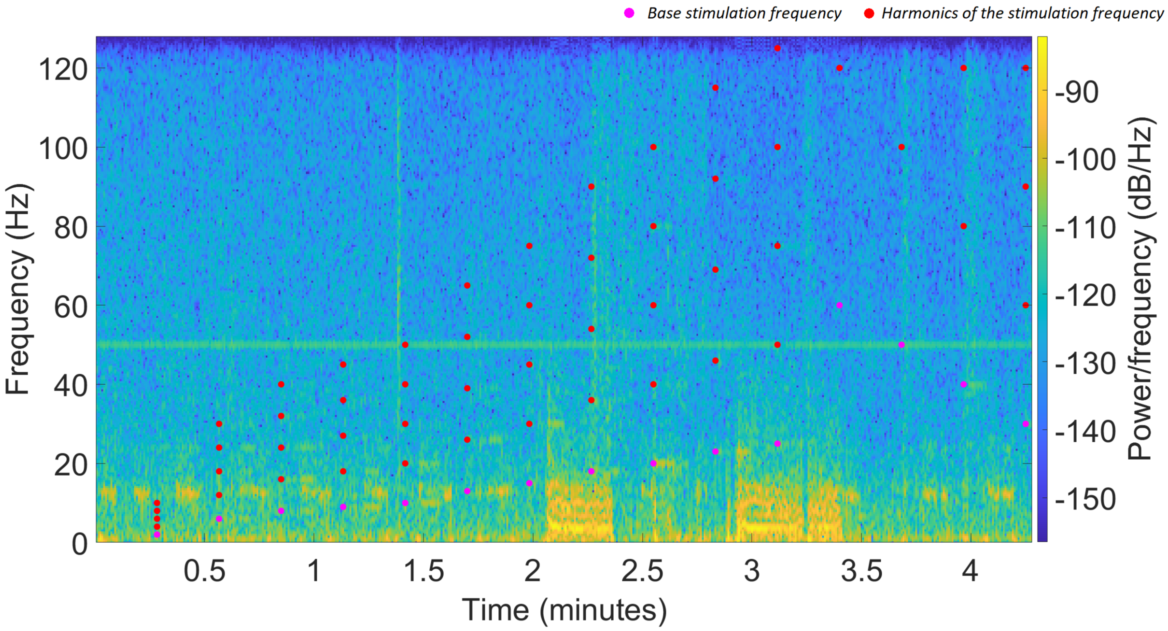


S2B: PARTICIPANT 14 (GGE NO PPR)

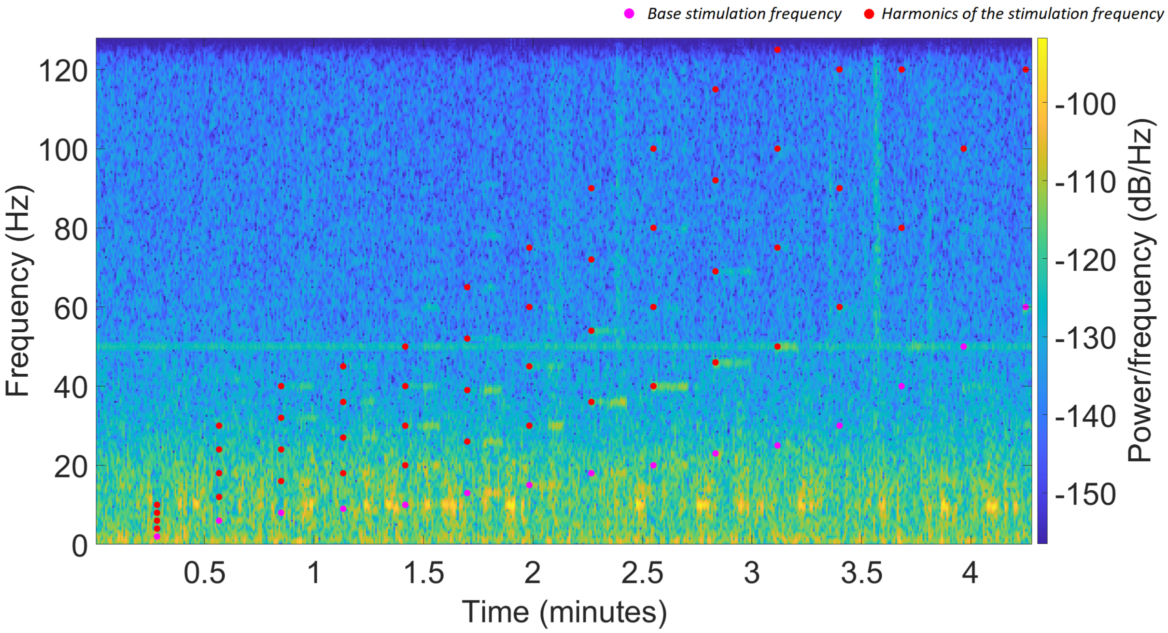


Figure S1: (cont...)

S2C: PARTICIPANT 28 (PNES)

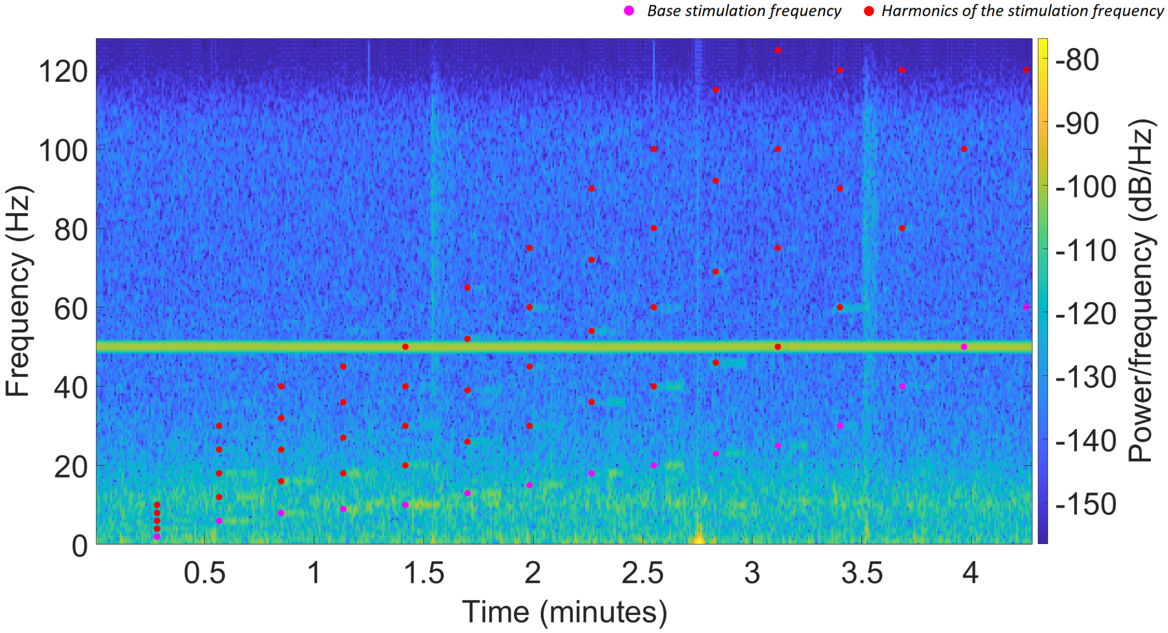


Figure S2: Spectrograms of participant 1 (subplot 2A), 14 (subplot 2B) and 28's (subplot 2C) stimulation trains, from the O1, O2, and O2 channels respectively. Spectrograms are calculated using MATLAB's spectrogram fnunction, with a 1s window, 'overlap' of a half second, and epoch length of 2s. Frequencies from 1-128Hz are shown along the y-axis, time in seconds is shown on the x-axis, and the power estimate for each frequency and time bin is shown as a colour described in each subplot's legend in power/frequency (dB/Hz), ranging from greatest power in yellow and least power in dark blue. For each plot, the frequency of stimulation is indicated as a pink filled dot at the start of each 10 second stimulation epoch, and each harmonic up to it's 5th multiple is shown as a red dot. In participant 1's spectrogram (subplot 1), their two PPR events are visible at approximately 2 minutes and 3 minutes as high power activity. High power lines at 50Hz are technological artifact.

S3A: PARTICIPANT 1

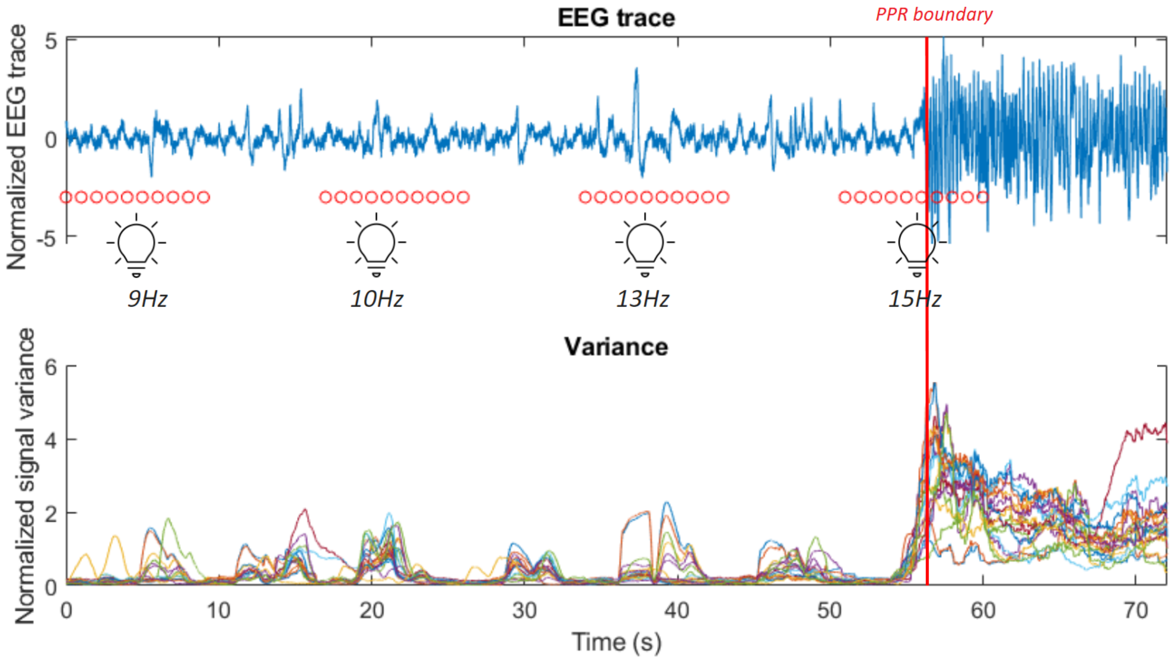


S3B: PARTICIPANT 2

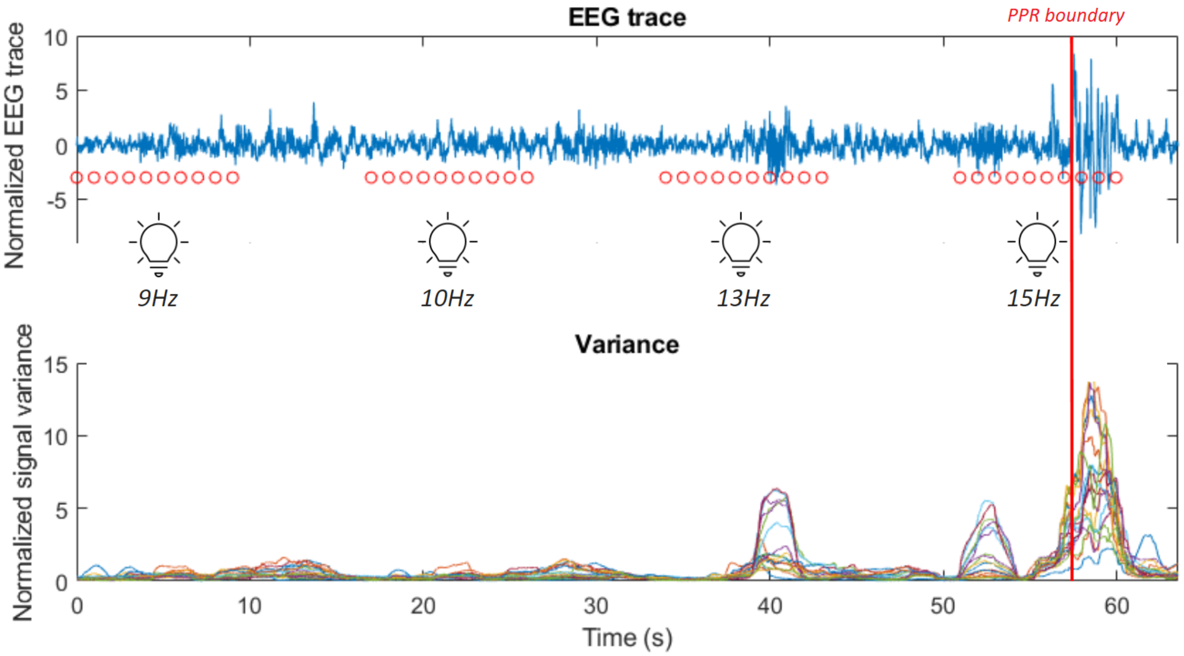


Figure S3: (cont...)

S3C: PARTICIPANT 8

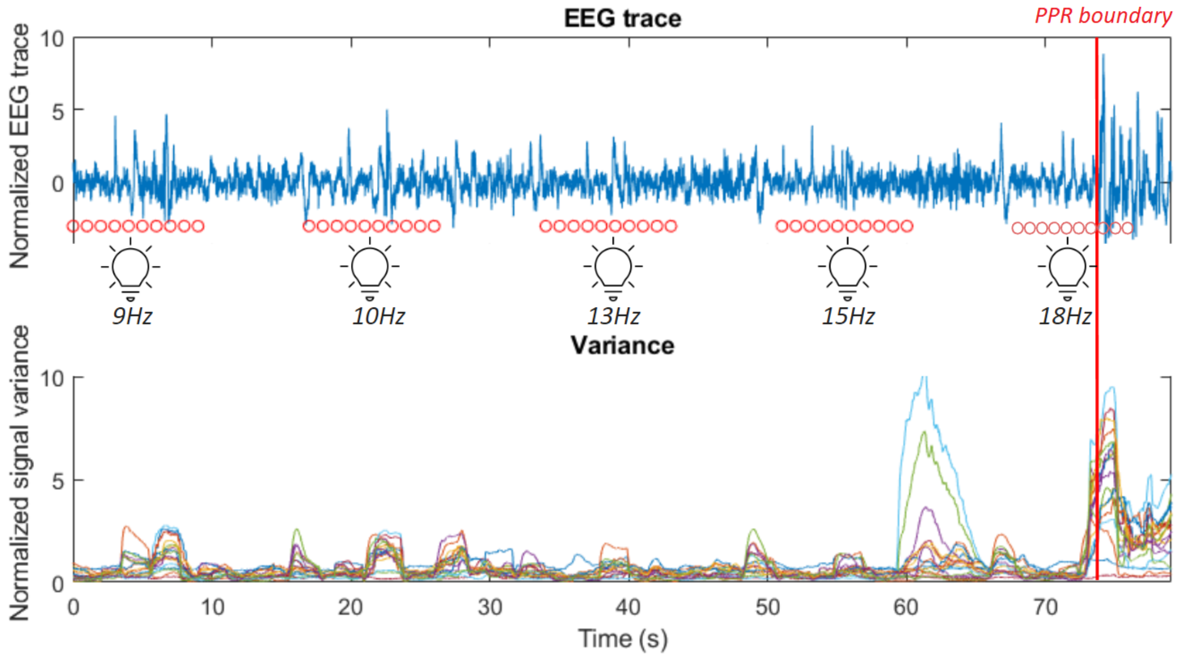


S3D: PARTICIPANT 10

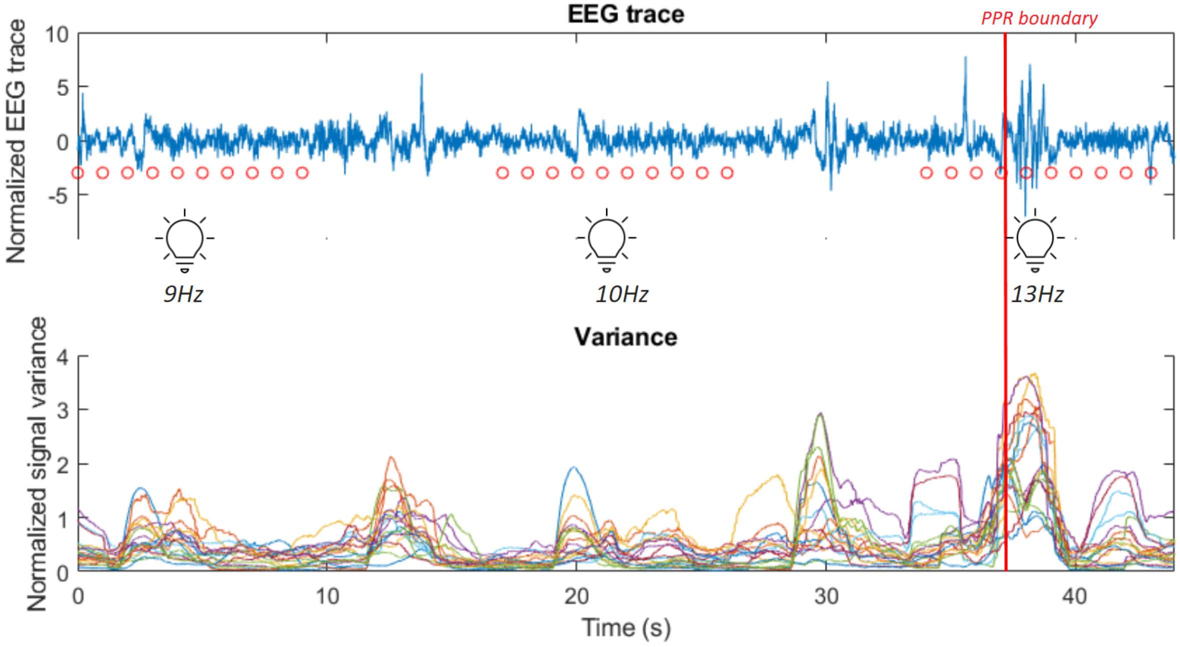


Figure S4: The first subplot shows the EEG trace (blue line) of participant 1 in the leadup to a PPR, visible as high amplitude spiking activity after the marked PPR onset boundary (vertical red line, defined as the onset of the PPR determined through a neurologist's inspection). Consecutive red dots mark 10 seconds of photic stimulation for the frequency (labelled below them). Below, the corresponding variance of the signal is displayed, using the shared time axis (y-axis), where different electrode channels are displayed as different coloured lines. The second, third, and fourth subplots displays the same information for participants 2, 8 and 10. Observe the individualised response to the same photic stimulation frequencies, though similar increased variance within tens of seconds of the boundary.

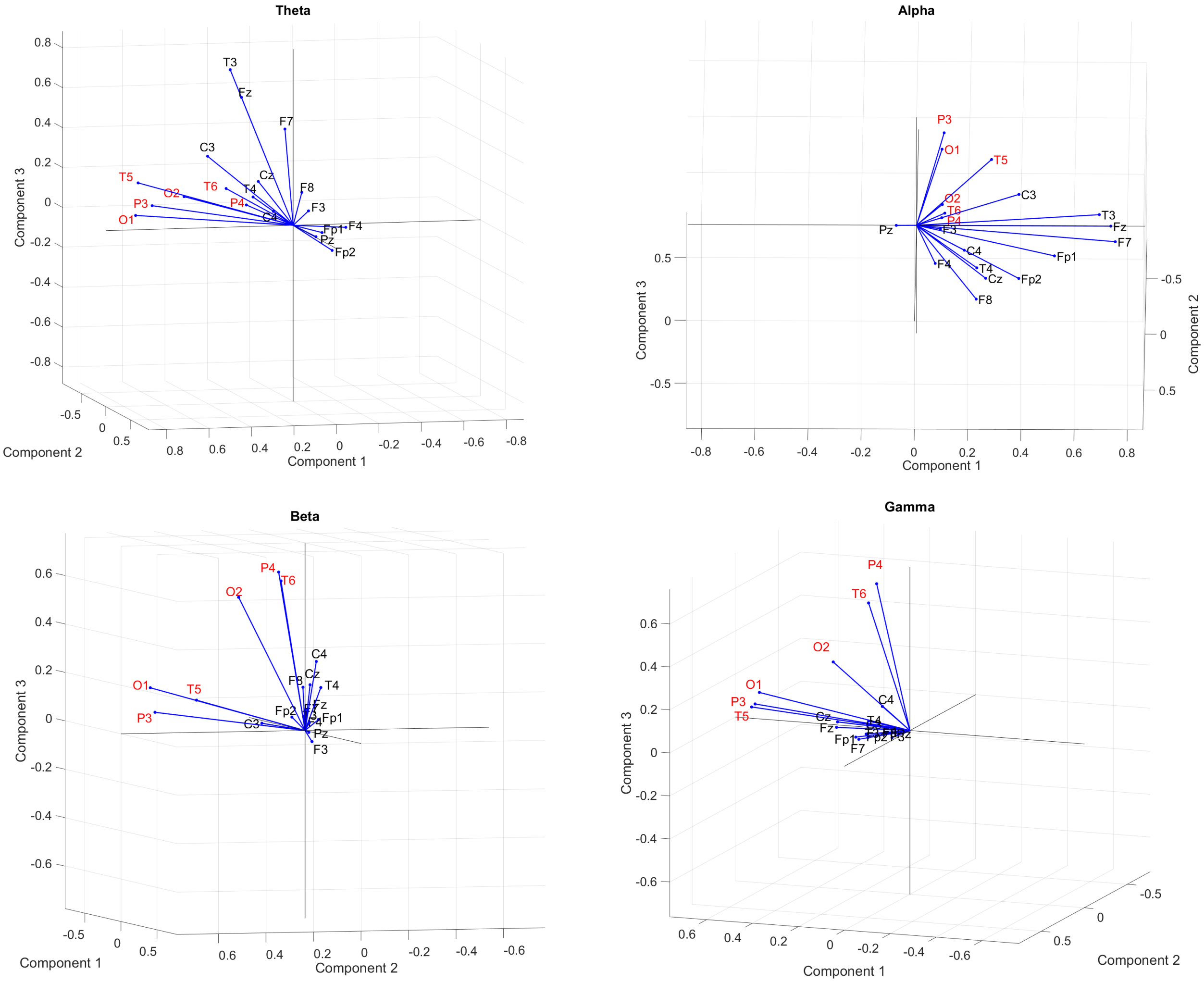


Figure S5: Factor loadings for each channel (blue vectors) plotted on the three dominant component axes, in different frequency bands: **(A)** Theta; **(B)** Alpha; **(C)** Beta; **(D)** Gamma. Posterior channels (P3, P4, T5, T6, O1, O2; labelled with red text) often show the largest variation, illustrated by the longer vector lengths. This suggests that this subset of channels demonstrated the greatest amount of change in their values.
